# NRN-EZ: an application to streamline biophysical modeling of synaptic integration using NEURON

**DOI:** 10.1101/2022.07.18.500394

**Authors:** Evan A. Cobb, Maurice A. Petroccione, Annalisa Scimemi

## Abstract

One of the fundamental goals in neuroscience is to determine how the brain processes information and ultimately controls the execution of complex behaviors. Over the past four decades, there has been a steady growth in our knowledge of the morphological and functional diversity of neurons, the building blocks of the brain. These cells clearly differ not only for their anatomy and ion channel distribution, but also for the type, strength, location and temporal pattern of activity of the many synaptic inputs they receive. Compartmental modeling programs like NEURON have become widely used in the neuroscience community to address a broad range of research questions, including how neurons integrate synaptic inputs and propagate information through complex neural networks. One of the main strengths of NEURON is its ability to incorporate user-defined information about the realistic morphology and biophysical properties of different cell types. Although the graphical user interface of the program can be used to run initial exploratory simulations, introducing a stochastic representation of synaptic weights, locations and activation times typically requires users to develop their own codes, a task that can be overwhelming for some beginner users. Here we describe NRN-EZ, an interactive application that allows users to specify complex patterns of synaptic input activity that can be integrated as part of NEURON simulations. Through its graphical user interface, NRN-EZ leverages the learning curve to run computational models in NEURON, for users that do not necessarily have a computer science background.

## Introduction

Information transfer within the brain relies on the ability of individual neurons to integrate synaptic inputs in space and time. In most instances, synaptic integration behaves as a non-linear system, where neither analytical solutions nor intuition can guide our understanding of the operating principles of neurons and circuits. To overcome this limitation, back in the ‘80s, scientists began to develop approaches to efficiently compute the Hodgkin Huxley branched cable equations [1-3]. These works laid the foundation for the development of realistic quantitative models through which investigators could determine how non-homogeneous ion channel distribution in neurons with complex architectures controls synaptic integration and action potential firing. The software NEURON was developed exactly for this purpose [1, 4-6]. To this day, its use continues to be instrumental to cross-validate experimental data, estimate experimentally inaccessible parameters and establish relationships between specific patterns of synaptic activity and firing output profiles. NEURON relies on an interpreted programming language based on a floating-point calculator with C-like syntax, called hoc (an acronym for high order calculator) [7]. Through hoc codes, users define specific parameters in the simulations, analyze data, calculate new variables, etc. An improved hoc interpreter was developed in 1985 by the Computing Science Research Center at Bell Labs but was not generally adopted by commercial Unix systems or Linux distributions, which instead rely on developments of earlier calculator languages like dc and bc. Newer versions of hoc continued to be implemented as part of the Plan 9 operating system by Bell labs in 2015 and considerable extensions have been made to run the current version of NEURON. Learning how to code in hoc is not an insurmountable task for a computer scientist, but can be intimidating for many biologists that have not received formal or informal training in programming. On the other hand, computational proficiency and literacy have become essential skills in a biologist’s toolbox, guiding data interpretation and experiment design. Therefore, it is important to make all these tools more accessible to the wider scientific community and junior trainees.

To overcome these hurdles, we developed an open source application called NRN-EZ, which is accessible to users with limited programming background. NRN-EZ allows users to: *(i)* load the 3D morphology of a digitally reconstructed neuron; *(ii)* add user-defined excitatory and inhibitory synaptic inputs; *(iii)* generate uniform, Poisson or any user-defined distribution for synaptic weight, location, and activation time of different types of synaptic inputs; and *(iv)* automatically generate hoc files that can then be compiled in NEURON. The NRN-EZ code is freely accessible, easy to use and can be modified by users as they see fit.

## Design and Implementation

### Software description

NRN-EZ is an open-source application that uses the Python programming language (version 3.6.9), the PyQt application (version 5.10.1) and the PyQtGraph module (version 0.11.0), build with PyInstaller 3.6. NRN-EZ was built on Ubuntu 18.04.5 LTS, Windows 10 and MacOS Big Sur 11.5.2. NRN-EZ uses a Model - View - Controller (MVC) software design, which divides the program logics into three interconnected elements through which users can generate one or more synaptic inputs to run compartmental models using NEURON.

The Model portion of NRN-EZ consists of the following classes: experiment (also referred to as session in the GUI), morphology, weight, location (which also handles the number of inputs) and timing. The morphology class loads a .swc file, generates an output graph shown in the left panel of the GUI, and creates a .nrn file. The weight, location and timing classes all do error checking on the user input data and they all handle generating data based on the user input. The experiment class (i.e., the session) does error checking, and generates .hoc files. Any data with Poisson and uniform distributions are created using Python’s built-in math module. These data persist in the memory while the application is running. These and any other user-defined input parameters are also saved when the user hits the “Run” button in the bottom left corner of the GUI. When saving the data, NRN-EZ “pickles” the data (pickling is a built in python module that saves python objects to flat files). Once saved, these .pkl files can be reloaded as a session into NRN-EZ at any time. The Model portion saves information into the following files: experimentClass.py, morphologyClass.py, locationClass.py, timeClass.py, weightClass.py. These files contain the class definitions for all the user input data. These also contain the output generation functions.

The View portion of NRN-EZ consists of the GUI built using PyQtGraph because of its ability to quickly generate large plots, like the one embedded in the NRN-EZ GUI. Accordingly, when users select a .swc file to load into NRN-EZ, the software quickly renders it graphically. The View portion saves information into the following files: gui.py, gui_helper.py. These files contain all the code for building the user interface.

The Controller portion of NRN-EZ transfers data from the GUI to the Model portion (e.g., when a user clicks the “Save” buttons in the middle and right portions of the GUI). In addition, the Controller portion transfers data from the Model to the GUI (e.g., when the user edits a module by clicking on its corresponding row in the tables or when the “Run” button is hit to execute the code and update the graph). The Controller portion saves information into the gui_handler.py files. This file contains the code to handle user actions.

The code also includes a file that checks for command line arguments (NRN-EZ.py), an error logger module (errorLogger.py), which generates the run log, error log, and debug log. It also handles generating the error messages. The configuration module (config.py) handles options that users can change without having to recreate and version the code (i.e the reference comment for the .hoc file outputs.). Last, the code stores information about the definitions of all global variables (globvar.py).

### Overview of the graphical user interface

NRN-EZ has a simple and intuitive graphical user interface (GUI; **Figure 1**), organized in three adjacent panels. Through the GUI, users can load the 3D structure of a biocytin filled and reconstructed cell (which is displayed as a 2D maximum intensity projection in the NRN-EZ interface) and populate it with inputs of varying weights, spatial distribution, and temporal patterns of activation. The 3D morphology of a cell can be generated *de novo* from 3D reconstructions of biocytin-fills using open source or proprietary software (e.g. SimpleNeuriteTracer, Bitplane Imaris, etc.). Alternatively, one can use 3D reconstructions available from publicly accessible repositories like NeuroMorpho.org (https://www.neuromorpho.org) [8, 9]. Each reconstruction is typically stored as a .swc file, an open source file format widely used in the neuroscience community. In each .swc file, cells are defined through a list of sub-cellular compartments, which users can populate with arbitrary passive and active conductances. A typical neuron contains four types of compartments: the soma, the axon, the apical and the basal dendrites. Each compartment is identified by its 3D Cartesian coordinates, its radius and parent compartment. The .mod file generated through NRN-EZ can then be imported in NEURON to analyze, for example, the spiking output of a neuron (see examples in **Figure 3-5**).

**Figure 1.**
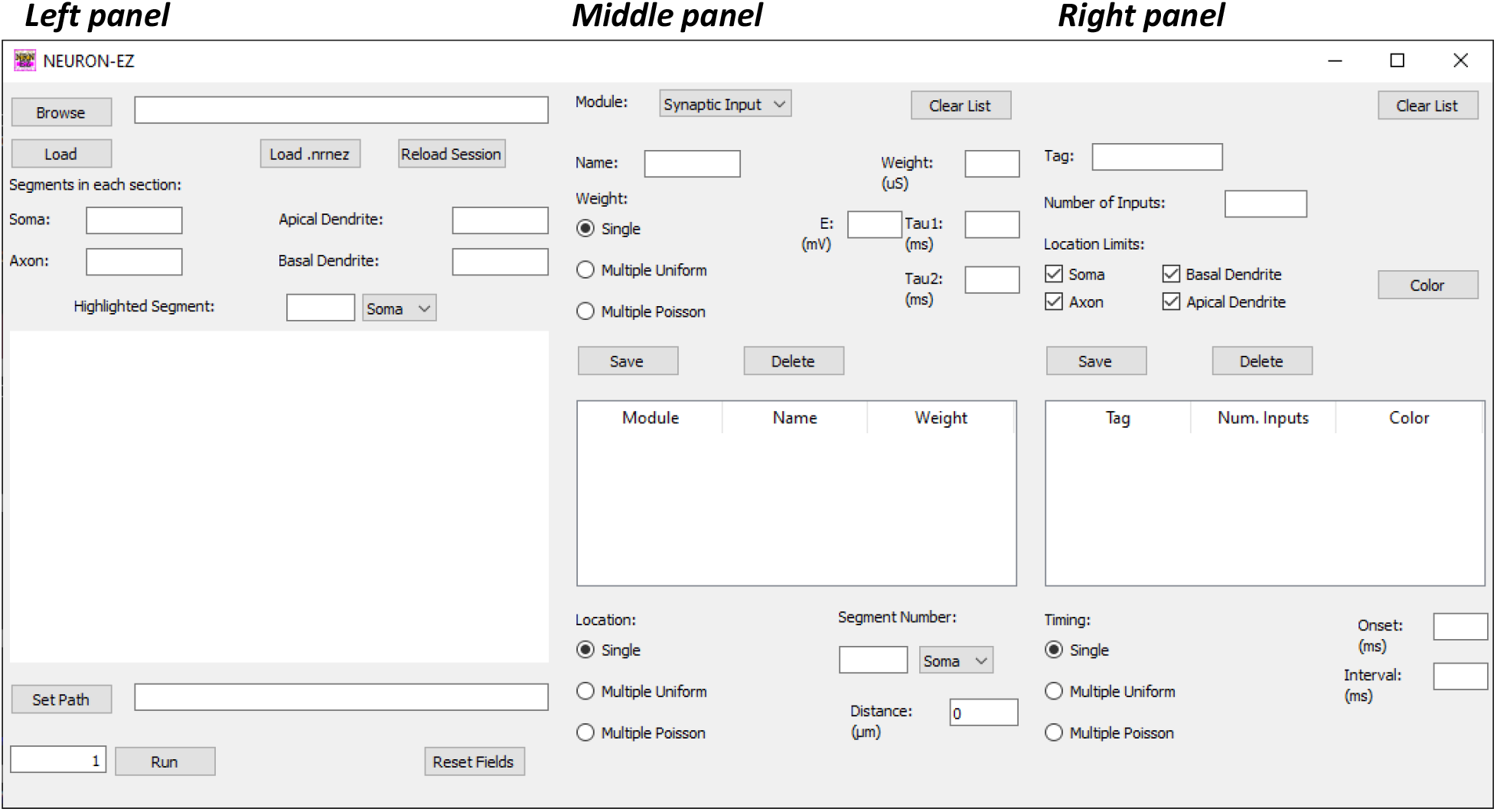
The NRN-EZ GUI. The GUI is composed of a left, middle, and right panel. The left panel allows users to upload a file containing information about the 3D neuron structure. Once the upload is finished, the users can see the number of segments in each of the four sections of the neuron. They can also pan through the neuron morphology, identify specific segments, and set the path to the directory, which will contain the output from NRN-EZ. The middle panel is the one through which users specify the type of module they plan to use: current steps or synaptic stimulations. They can use multiple modules and assign them an arbitrary name. The weight, kinetics and timing of activation can be selected through the parameters described in the upper portion of the panel. The bottom portion allows users to select the location and spatial distribution of these inputs. The right panel allows selecting the number of inputs and the limits for their spatial location. For example, one may decide to confine synaptic inputs of a given type only to one dendritic compartment. NRN-EZ allows selecting multiple types of inputs, with unique locations and weights that can be integrated to run NEURON simulations of synaptic integration.

**Figure 2.**
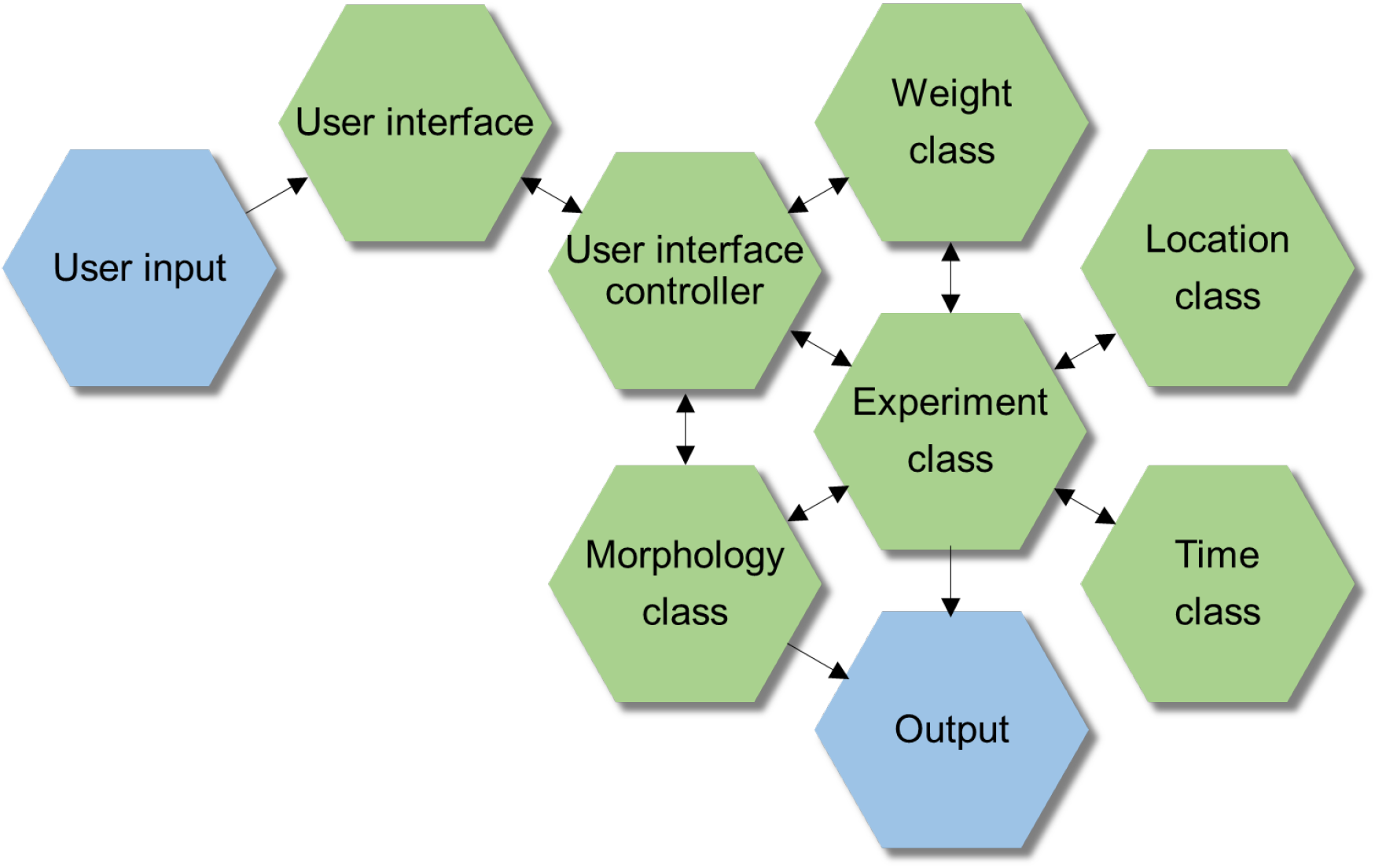
Software architecture of NRN-EZ. Schematic representation of the fundamental structure and design for the NRN-EZ software. The blue hexagons highlight the start and end points of the logical workflow (i.e., the parameter choice from the GUI and the output file generated using NRN-EZ. The green hexagons identify intermediate workflow steps.

### The left pane of the GUI: visualizing cells and setting the path for the working environment

The first step when using NRN-EZ is to either: *(i)* load a .swc file, *(ii)* reload a .pkl file, which allows users to recreate the morphology and reload all input parameters (e.g., synaptic weight, onset, etc.); or *(iii)* load a .nrnez file, which allows to load some default values into the GUI, including values for different input parameters. The .swc file can be loaded by first navigating through its location folder using the “Browse” button in the top left corner of the GUI, and by clicking the “Load” button below it. The entire NRN-EZ session, which contains the stimulation parameters, and the cell morphology can be loaded by clicking the “Reload Session” button. Users can press the “Load .nrnez” button to upload a set of default parameters that they find convenient to have as part of their default screen. A sample .nrnez file is included in the NRN-EZ GitHub repository.

Once the .swc file is uploaded, one can visualize the reconstructed cell. The axis units change from µm to mm depending on the size of the cell. The GUI displays the segment composition of the cell as a list that includes the number of segments in the soma, axon, apical and basal dendrites sections. In the interactive 2D Cartesian plane embedded in the GUI, these sections appear as color-coded in black (soma), magenta (axon), blue (apical dendrites) and cyan (basal dendrites). By right clicking and dragging the mouse up and down, we can zoom in and out the Cartesian reference system along the y-axis. By right clicking and dragging the mouse right and left, we can zoom in and out along the *x*-axis. The mouse scroll wheel can be used to zoom in simultaneously along the *x,y*-directions. By clicking and dragging the left mouse button, we can move the neurons around, in any direction. The identity of a segment can be revealed in the fields above the graph by drawing a line across it. The selected segment is highlighted with its complementary color. Alternatively, users can highlight specific segments by typing their segment identity number and segment identifier in the corresponding fields above the graph area (e.g., by typing “100” and “Apical”).

The left portion of the GUI also allows selecting the destination folder for the output files generated by NRN-EZ, which includes information about the weight, location and activation time of various synaptic inputs, or the amplitude and duration of current step injections. This can be done by navigating to the destination folder using the “Set Path” button, and then clicking the ”Run” button to execute the code specified through all other panels in the GUI. Users can select to run the code one or multiple times by typing a number in the field on the right-hand side of the “Run” button. To clear all parameters from the whole GUI, one can use the “Reset Fields” button on the right-hand side of the “Run” button.

### The middle panel of the GUI: selecting modules and setting their weight properties selection

In the middle panel of the GUI, users can select the type of stimulus (or module) to be used in their simulations. There are three main types of modules that can be used and that can be distributed across one or more segments. The “I-Step” module allows injecting current steps of any given amplitude and duration into a specified segment of a given compartment. The second module, called “Synaptic Input” is used to generate synaptic inputs based on a two-state kinetic scheme synapse described by rise time and decay time constants tau1 (rise) and tau2 (decay), respectively (see NEURON documentation). The name of each “I-Step” and “Synaptic Input” module type can be set arbitrarily by the user (e.g., AMPA, NMDA, step) in the box called “Name”, a useful feature to prevent potential nomenclature ambiguities when using multiple types of current steps or synaptic inputs in the same simulation. The third module type, “Custom”, allows the use of point processes (i.e. voltage or ligand-gated ion channels) with custom properties defined using .mod files. When using the “Custom” module type, the “.mod File” button prompts users to navigate to the .mod file to be used. In this case, once the .mod file is selected, NRN-EZ automatically sets the name the module in the table to match the name of the .mod file. Once the module type name has been assigned, the users can select whether the weight of each input should remain the same (“Single”) or vary according to a uniform (“Multiple Uniform”) or Poisson distribution (“Multiple Poisson”). If the “Multiple Uniform” or “Multiple Poisson” options are selected, then the GUI prompts the user to select the mean and standard deviation of the current (“I-Step”) or of the synaptic weight (“Synaptic Input” and “Custom”). When using the “Synaptic Input” module type, the users also need to select the reversal potential (“E”), and the rise and decay time constants (tau1 and tau2, respectively). Once these parameters have been set, they are saved together with the module type and name and are displayed in the table at the center of this panel. The parameters of any saved module type can be changed by clicking on their corresponding row in the table, and they can be stored by clicking the “Save” button. The “Delete” button can be used to delete any current step or synaptic input that is no longer needed.

The bottom portion of the panel can be used to set the location and spatial distribution of each module type. The options for the spatial distribution are analogous to those mentioned when describing the synaptic weight. Accordingly, any given module type can be delivered to a single location identified by its segment number (“Single”) or to multiple locations according to either a uniform (“Multiple Uniform”) or Poisson distribution (“Multiple Poisson”). The mean of their spatial distribution is calculated using Euclidean distance from the user-selected segment.

### The right panel: stimulus tags, saving tags, and input timing

The right panel is used to label, or tag, a group of module types. Modules with the same tag are applied to the same exact locations and are activated at the same time. For example, assigning the same tag to both AMPA and NMDA module types can be used to create synapses that contain both AMPA and NMDA receptors at the same exact location. The user can select how many of these inputs should be applied to their cell of interest. The location of stimuli labelled by the same tag can be confined to one or more specific domains of the cell, for example the basal and/or apical dendrites. One can use a color-coded scheme for each tag. NRN-EZ will generate random distributions for stimulus magnitude, location and timing, independently for each stimulus tag. Users can decide the temporal pattern of activation of inputs with a given tag. These inputs can be delivered simultaneously by selecting “Single” and setting the variable “Interval” to zero. They can also be delivered with a constant time interval between each other (“Single”) or with a uniform (“Multiple Uniform”) or Poisson temporal distribution (“Multiple Poisson”), each defined by a mean time interval and standard deviation. The “Onset” field is used to set a delay between the beginning of the simulation and the activation time of the first input. “Interval” defines the time interval between consecutive stimuli. The table of saved tags in the right panel functions similarly to the saved stimuli table in the middle panel, where users can delete or edit the parameters assigned to a tag by selecting its corresponding row in the table.

### Generating the inputs and running the simulation

Once all the desired stimulus tags have been created, the code can be run by clicking the “Run” button in the bottom left corner of the GUI. The After clicking the “Run” button, NRN-EZ displays each input in the Cartesian coordinate system of the left panel as a dot with a different color for each tag. The output files of NRN-EZ (.mod, .hoc, .nrn, .pkl, .swc,) are stored in a folder named with the date and time of its creation, in the destination folder identified by the “Set Path” button (left panel). The folder created by NRNR-EZ contains a sub-folder with a .hoc file with the synaptic weight information and multiple .dat files with input location, time and weight information. In each .dat file, the first row contains two numbers, corresponding to the number of rows and columns in the file. All these files are used to run a Hodgkin-Huxley simulation using NEURON.

In order to run a NEURON simulation, one should first compile the files located in the NRN-EZ output folder. This is done by running mknrndll (in the NEURON software), navigating to the output folder of NRN-EZ, and clicking the “Make nrnmech.dll” button (in the NEURON software). After compiling the files, the user can run the nrnez.hoc file, created by NRN-EZ and stored in its the output folder, by using NEURON. This will create a .ses (session) NEURON file containing all the parameters specified for the simulation, which can be changed from the NEURON GUI (e.g. simulation length).

## Results

We tested and validated NRN-EZ by using the 3D morphology of a CA1 pyramidal cell in the mouse hippocampus, obtained using biocytin fills and confocal laser scanning microscopy image acquisition (NeuroMorpho NMO_139428) [10]. We assigned this neuron a resting membrane potential of -65 mV, and the passive and active membrane properties of a CA1 pyramidal neuron described previously [10]. Briefly, the model included voltage-gated sodium channels (Na_V_), A-type potassium channels (K_A_), delayed-rectifier potassium channels (K_DR_), calcium-dependent potassium channels (K_Ca_), muscarinic receptor activated potassium channels (K_M_), hyperpolarization activated cationic channels (I_h_), and L, N, T-type calcium channels. All simulations were performed using a variable time step and the code for the NRN-EZ simulations is deposited on GitHub (https://github.com/scimemia/NRN-EZ). The NEURON code is shared on the ModelDB database (Acc n. 267419).

In a first series of simulations, we used NRN-EZ to change the location of a single excitatory input, with a conductance of 1 nS, from 100 µm (segment #3285) to 200 µm (segment #943) and 300 µm away from the soma (segment #2420; **Figure 3**). Using the middle panel of the NRN-EZ GUI, we set the reversal potential for AMPA EPSPs at E=0 mV. Each AMPA input had a rise time constant tau1=0.7 ms and a decay time constant tau2=2 ms. We selected “Single” location and entered the corresponding segment number and type (e.g., 3285 Apical) and pressed Save. In the right panel, we tagged this input as either Proximal, Medial or Distal and set the number of inputs to 1. The location limit was set to Apical Dendrite, to match the location of the selected segment. We selected red as the color for visualizing the input (**Figure 3A**). At the bottom of this panel, for timing, we selected “Single”, set the Onset to 200 ms, clicked the Save button and then the Run button in the left panel. The results of the NEURON simulations performed using these files are shown in **Figure 3B**. The amplitude of the somatic EPSP decreased as the input location moved further away from the soma, while its kinetics (rise and decay time) became slower. These findings are consistent with the basic principles of cable theory, used to describe the physiological properties of passive and active input propagation along dendrites [11]. If rather than using a single input, we used 100 of them, and distributed them with a Poisson location centered 100-300 µm away from the soma, we recorded a single action potential at the soma, arising on top of an EPSP that became less prominent for more distal inputs (**Figure 3C-D**). When the excitatory inputs were not synchronous, but had a Poisson distribution for their activation time, the cell produced multiple action potentials if their location was <200 µm away from the soma (**Figure 3E**). A train of action potentials could be generated by varying the synaptic weight using a Poisson distribution (**Figure 3F**). This is important, as it suggests that varying the strength and timing of the excitatory inputs increases the firing output probability of CA1 pyramidal cells, and therefore the temporal fidelity with which they relay information to the entorhinal cortex. The frequency of the trains decreased as the inputs became more distal (**Figure 3F**). We detected a qualitatively similar trend when using a uniform rather than a Poisson distribution for synaptic input timing and weight. However, in this case, the firing output of the cell decreased more abruptly at increasing distances from the soma (**Figure 3G-H**).

**Figure 3.**
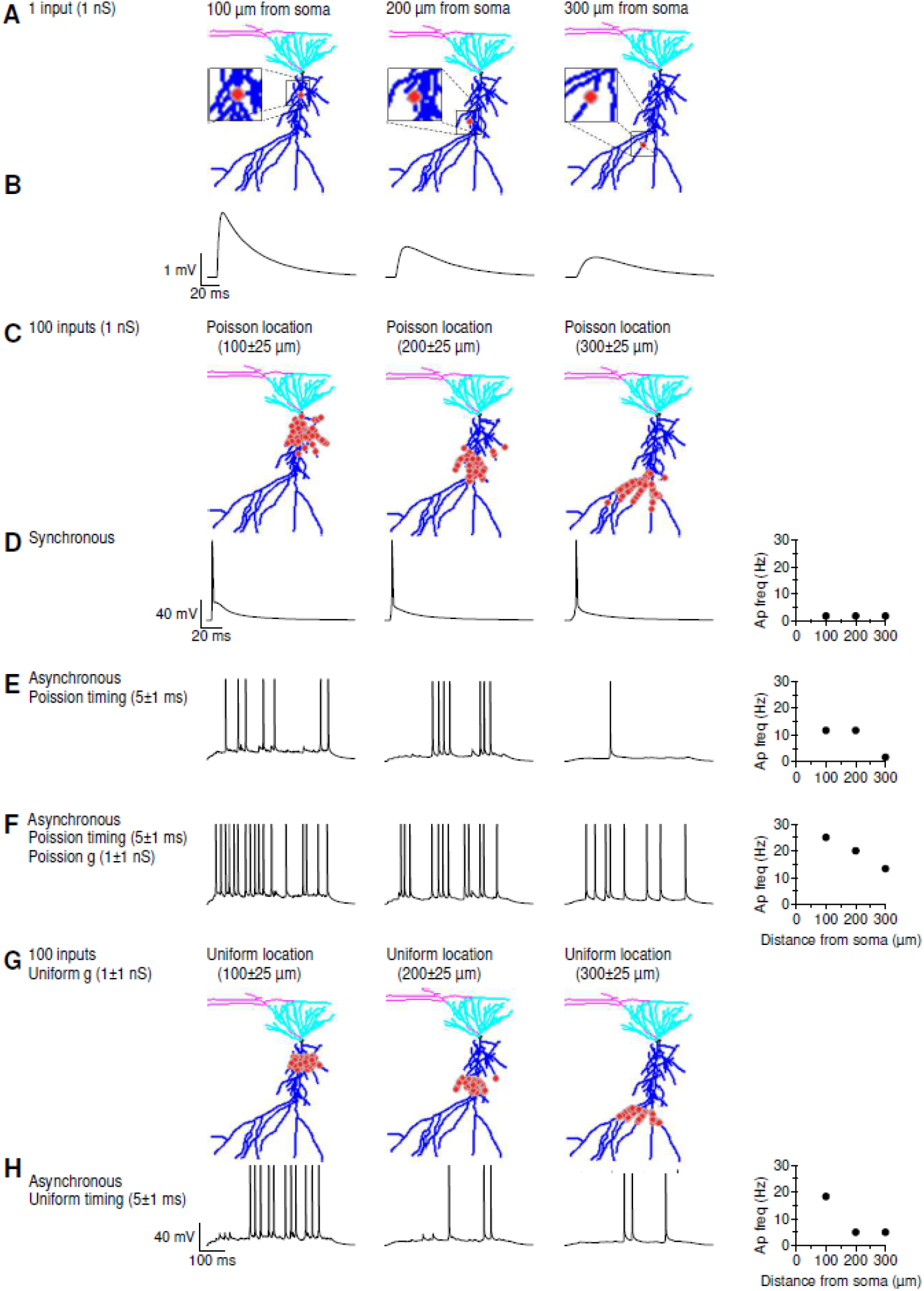
Sample Application of NRN-EZ for summation of supra-threshold EPSPs. **(A)** We used NRN-EZ to place one AMPA excitatory input at 100-300 µm from the soma of a 3D reconstructed, biocytin-filled CA1-PCs. The maximum intensity projection of the CA1-PC includes the soma (*black*), basal dendrites (*cyan*), apical dendrites (*blue*) and the location of the input (*red*). The synaptic weight was set to 1 nS. **(B)** Somatic recording obtained using the NEURON simulation environment. The action potential frequency recorded at the soma for inputs located at increasing distances is plotted in the right panel. **(C)** As in A, for 100 inputs with a Poisson placement along the distal dendrite of the CA1-PC. **(D)** As in B, in response to the synchronous activation of the 100 inputs described in C. **(E)** Somatic membrane potential recorded in response to the asynchronous activation of the 100 excitatory inputs described in C. **(F)** As in E, for synaptic inputs with a Poisson conductance distribution. **(G)** As in C, for 100 inputs with a uniform spatial distribution centered 100-300 µm away from the soma. **(H)** As in F, for 100 inputs with uniform spatial distribution and uniform timing of activation.

To understand how variability in the location, strength, and timing of activation of different synaptic inputs affect the integration of sub-threshold inputs, we repeated the simulations using a smaller conductance for the AMPA input (7.5 pS; **Figure 4**). In this case, we quantified the maximal depolarization at the soma and the time integral of the change in membrane potential (i.e., the area under the curve, AUC). Consistent with the data previously described for **Figure 3A-B**, a single excitatory input generated a smaller and slower EPSP as it moved from proximal to distal locations (**Figure 4A-B**). This general principle held true also when analyzing the somatic output in response to 100 synchronous (**Figure 4C-D**) and asynchronous inputs (**Figure 4E-F**). The magnitude of the somatic depolarization, however, was larger when the value of the synaptic weight varied according to a Poisson distribution (**Figure 4F**). This is consistent with the results described for supra-threshold stimuli (cf. **Figure 4E-F** and **Figure 3E-F**). This effect is less pronounced when the variability of synaptic weights is described through a uniform, rather than a Poisson, distribution (**Figure 4G-H**).

**Figure 4.**
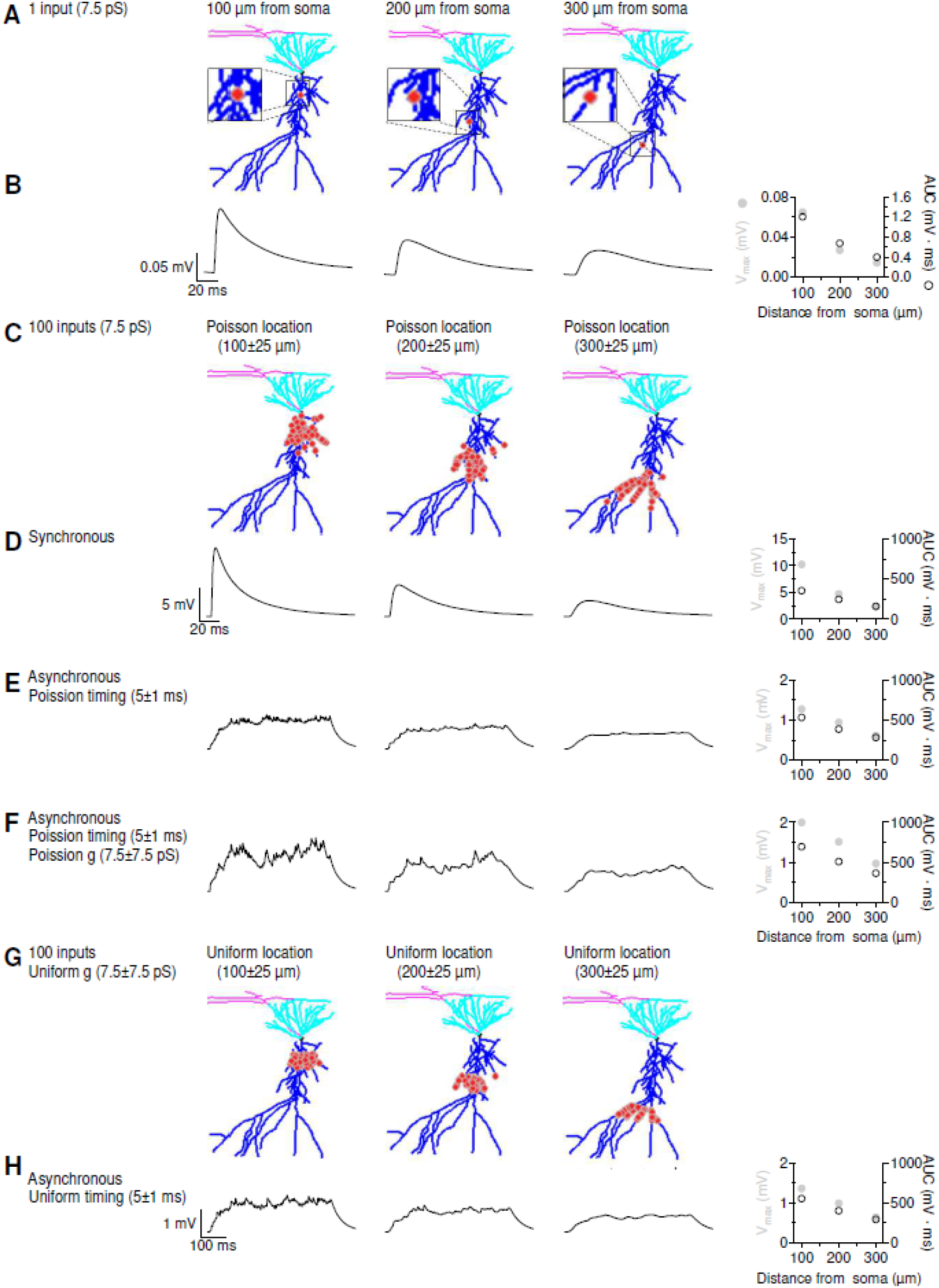
Sample Application of NRN-EZ for summation of sub-threshold EPSPs. **(A)** We used NRN-EZ to place one AMPA excitatory input at 100-300 µm from the soma of a 3D reconstructed, biocytin-filled CA1-PCs. The maximum intensity projection of the CA1-PC includes the soma (*black*), basal dendrites (*cyan*), apical dendrites (*blue*) and the location of the input (*red*). The synaptic weight was set to 7.5 pS. **(B)** Somatic recording obtained using the NEURON simulation environment. The maximum depolarization and the area under the curve (AUC) of the somatic EPSP for increasing distances from the soma are plotted in the right panel. **(C)** As in A, for 100 inputs with a Poisson placement along the distal dendrite of the CA1-PC. **(D)** As in B, in response to the synchronous activation of the 100 inputs described in C. **(E)** Somatic EPSPs recorded in response to the asynchronous activation of the 100 excitatory inputs described in C. **(F)** As in E, for synaptic inputs with a Poisson conductance distribution. **(G)** As in C, for 100 inputs with a uniform spatial distribution centered 100-300 µm away from the soma. **(H)** As in F, for 100 inputs with uniform spatial distribution and uniform timing of activation.

The GABA_A_ component had a rise time constant tau1=0.5 ms and a decay time constant tau2=7 ms. The reversal for GABAergic synaptic inhibition was E=-75 mV, meaning that its driving force was substantially smaller than that of glutamatergic excitation. Therefore, the general principle of distance-dependence for the attenuation of synaptic inputs were smaller (though qualitatively similar) compared to those described for sub-threshold excitation. (cf. **Figure 4** and **Figure 5**). Even in this case, the Poisson distribution of synaptic weights was more effective than the uniform distribution at hyperpolarizing the somatic membrane potential.

**Figure 5.**
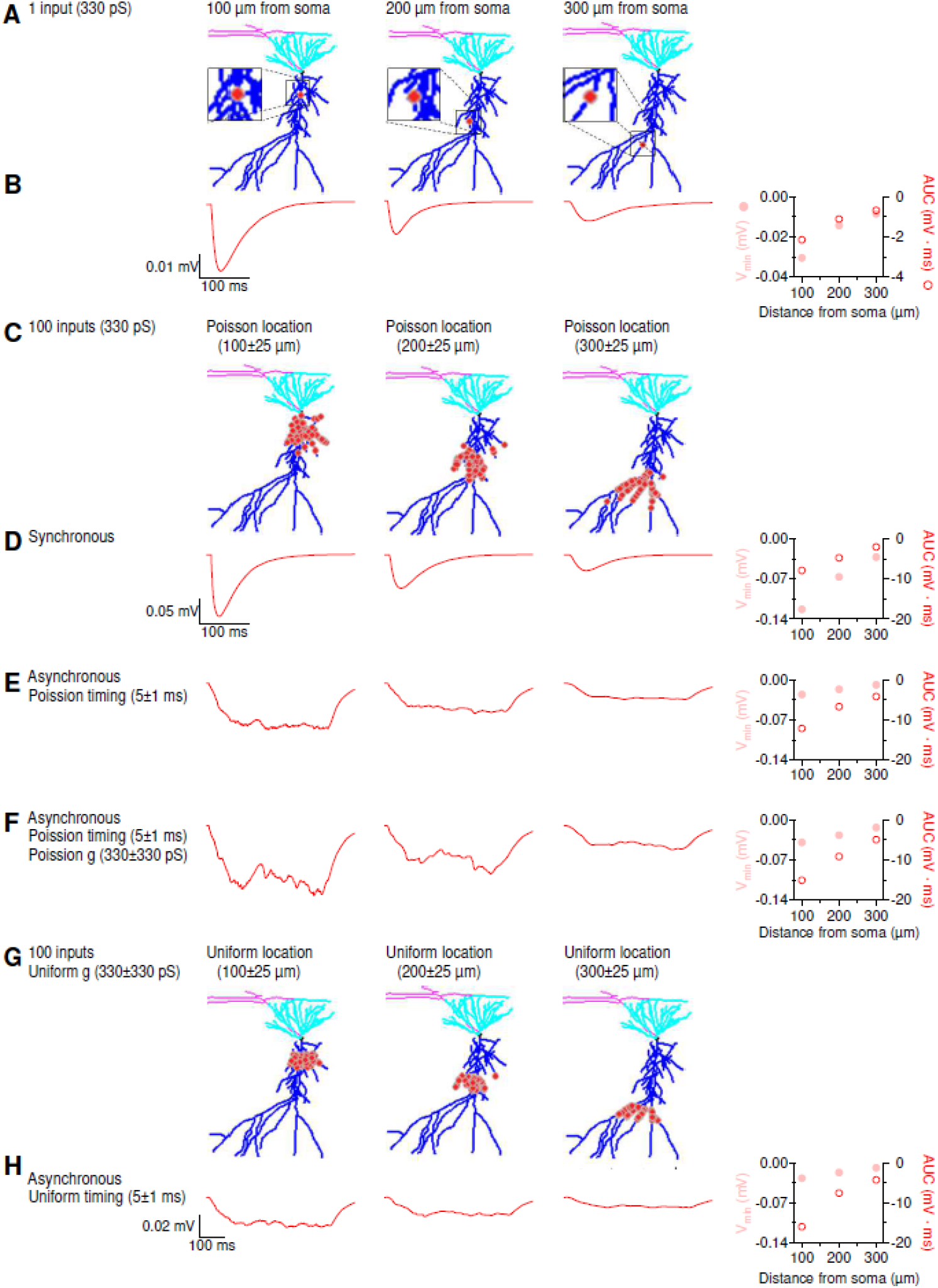
Sample Application of NRN-EZ for summation of IPSPs. **(A)** We used NRN-EZ to place one GABA_A_ inhibitory input at 100-300 µm from the soma of a 3D reconstructed, biocytin-filled CA1-PCs. The maximum intensity projection of the CA1-PC includes the soma (*black*), basal dendrites (*cyan*), apical dendrites (*blue*) and the location of the input (*red*). The synaptic weight was set to 330 pS. **(B)** Somatic recording obtained using the NEURON simulation environment. The minimum value of the membrane potential (V_min_) and the area under the curve (AUC) of the somatic IPSP for increasing distances from the soma are plotted in the right panel. **(C)** As in A, for 100 inputs with a Poisson placement along the distal dendrite of the CA1-PC. **(D)** As in B, in response to the synchronous activation of the 100 inputs described in C. **(E)** Somatic IPSPs recorded in response to the asynchronous activation of the 100 inhibitory inputs described in C. **(F)** As in E, for synaptic inputs with a Poisson conductance distribution. **(G)** As in C, for 100 inputs with a uniform spatial distribution centered 100-300 µm away from the soma. **(H)** As in F, for 100 inputs with uniform spatial distribution and uniform timing of activation.

Together, these results confirm some fundamental principles of cable theory but also allow exploring new parameters related to the variability in the spatial and temporal properties of synaptic integration in central neurons.

## Availability and Future Directions

In this work, we describe an application, NRN-EZ, which streamlines hoc file generation for NEURON compartmental models. The software and documentation for NRN-EZ are freely available for download from the Scimemi Lab website (https://sites.google.com/site/scimemilab2013/software) and GitHub repository (https://github.com/scimemia/NRN-EZ). Detailed instructions on software installation and operation are available from the GitHub repository, which also includes sample .swc files of 3D digitally reconstructed neuron morphologies. NRN-EZ can be downloaded as a standalone application for Windows, Mac OS, and Linux.

An important feature of NEURON, a simulation environment to run compartmental models of individual neurons and networks, is that it allows users to test how neuronal architecture, ion channel and synaptic input distribution and timing of activation affect firing output and computational efficiency [12]. Since its first development, NEURON has been used in more than 2,000 publications focusing on topics that range from integration, basic mechanisms of synaptic transmission, dendritic spike initiation, to neuron excitability, etc. (see https://neuron.yale.edu/neuron/publications/neuron-bibliography). NEURON incorporates a programming language based on hoc, through which users can write codes to define the weight and kinetics, spatial distribution and activation time of synaptic inputs or of points for current injections. Over the years, NEURON has become established in the neuroscience community as a powerful environment capable of handling complex neuron geometry and biophysical mechanisms, essential to prompt and constrain new hypothesis on neuronal function. It has indirectly contributed to the establishment of major scientific enterprises, like the Human Brain Project, Active Brain Mapping and Human Connectome Project, which aim to find the connection between the cellular principles of neuronal coding and cognitive/motor performance [13-19]. For many investigators with limited computer science knowledge, as well as undergraduate and graduate trainees that do not major in computer science but remain interested in understanding information processing in the brain, the learning barrier for NEURON can be high. This is particularly true when attempting to simulate complex activation patterns for different types of synapses with unique spatial distributions.

NRN-EZ is designed to leverage this barrier, facilitating the implementation of complex compartmental models by users with limited computer science experience. Being an open-source code, NRN-EZ can be implemented by anyone, which will be particularly useful for researchers around the world. NRN-EZ is flexible to cover a broad diversity of simulation scenarios, from current step injections to synaptic input activation. It allows to simulate synapses with mixed receptor populations, like glutamatergic synapses with AMPA and NMDA receptors.

We aim to keep NRN-EZ as simple, versatile, and useful as possible to make it a perfect entry tool to facilitate NEURON computations. Being an open source tool will allow users around the world to contribute to future implementation, which can include, for example, the implementation of an interface to write hoc codes for neural network simulations.

## Acknowledgments

This work was funded by the SUNY Albany Research Foundation NSF IOS-1655365 and IOS-2011998 to A.S. The funders had no role in study design, data collection and analysis, decision to publish, or preparation of the manuscript.

## Author Contributions

Conceptualization: EAC, AS

Data curation: AS

Formal analysis: MAP

Funding acquisition: AS

Investigation: AS

Methodology: EAC

Project administration: AS

Resources: AS

Software: EAC

Supervision: AS

Validation: MAP, AS

Visualization: EAC

Writing – original draft: AS

Writing – review & editing: MAP, AS

